# A-TWAS: An aggregated transcriptome-wide association study model incorporating multiple Bayesian priors

**DOI:** 10.1101/2025.01.27.635054

**Authors:** Yilan Liang, Han Wang, Yan Dora Zhang

**Author notes:** Co-first authors (Yilan Liang, Han Wang).

## Abstract

**Motivation:** Transcriptome-wide association study (TWAS) is a significant methodology utilized for identifying associations between genes and diseases by integrating transcriptome and genome-wide association studies (GWAS) data. The approach has been successful in pinpointing risk genes for various diseases, including Alzheimer’s disease, schizophrenia, and different types of cancers. TWAS typically involves two key steps: imputation and association analysis. In the imputation step, the original PrediXcan employs the elastic-net model, while subsequent research endeavors have delved into more intricate models such as FUSION and TIGAR. Despite these advancements, the existing individual methods may not capture the intricate genotype-expression relationships in a comprehensive way. Given the complexity of genetic contributions of genotypes to gene expression, sophisticated modeling techniques are imperative. In response to this, we have introduced Aggregated-TWAS (A-TWAS), a comprehensive tool that amalgamates multiple imputation models to accommodate diverse genetic architectures. A-TWAS utilizes several continuous shrinkage priors, including Laplace, Horseshoe, and Horseshoe+, to ensure precise imputation results while incorporating the benefits of Bayesian variable selection methods. In the association phase of TWAS, we employ the aggregated Cauchy association test (ACAT) to obtain an omnibus *p*-value.

**Result:** We demonstrate the effectiveness of A-TWAS through comprehensive simulations and real data analyses. Simulation studies highlight that A-TWAS produces substantial improvements in predictive *R*^2^ and statistical power, while maintaining the type error I at a low level. Furthermore, TWASs are conducted on schizophrenia and obsessive-compulsive symptoms datasets. In comparison to standard methods, the Bayesian imputation model demonstrates superior accuracy, and A-TWAS, by integrating multiple Bayesian priors, identifies a notably greater number of disease-relevant genes.

**Availability and Implementation:** The source code of A-TWAS is available at https://github.com/Yilan-Liang/A-TWAS.

## 1 Introduction

Genome-wide association studies (GWAS) have identified tens of thousands of genetic variants associated with complex traits and diseases, offering unprecedented insights into genetic architectures and disease mechanisms [23]. However, GWAS results primarily identify associations at the variant level, which complicates biological interpretation and limits their utility in understanding gene-level mechanisms. Transcriptome-wide association studies (TWAS) have emerged as a powerful tool to bridge this gap by integrating transcriptomic and GWAS data to identify gene-trait associations [11]. TWAS leverages gene expression as an intermediate phenotype, linking genotype to disease phenotypes through imputed gene expression, and has been successful in identifying risk genes for numerous diseases, such as Alzheimer’s disease [22, 14], schizophrenia [13], and multiple types of cancers [17, 24, 5, 6].

The implementation of TWAS involves two main stages [11]. The first stage trains predictive models to impute gene expression based on genotype data, while the second stage tests for associations between imputed gene expression and phenotypes using GWAS summary statistics. Despite its utility, a key challenge in TWAS lies in accurately modeling the complex relationship between genotype and gene expression. This relationship is influenced by various factors, including cis-regulatory elements, tissue specificity, and disease-specific mechanisms, making it highly variable across tissues and traits. Existing models often assume simplified or fixed structures for this relationship, which may fail to comprehensively capture the underlying genetic architecture of gene expression. This limitation motivates the development of more flexible and robust approaches to improve imputation accuracy and downstream gene-trait association detection.

Classic TWAS methods, such as PrediXcan [10], utilized elastic-net regression to model the cis-regulatory effects of single nucleotide polymorphisms (SNPs) on gene expression. Subsequent methods have employed more sophisticated statistical frameworks to improve imputation accuracy. For example, FUSION [12] uses Bayesian Sparse Linear Mixed Models (BSLMM) to capture both sparse and polygenic genetic effects, while TIGAR [18] adopts a nonparametric Bayesian model to flexibly model local expression quantitative trait expression quantitative trait loci (cis-eQTL) effect sizes. Despite these advancements, single-model approaches are limited by their reliance on specific model assumptions, which may not align with the true underlying genetic structure of gene expression. This misalignment can result in suboptimal imputation performance, particularly when the genetic architecture varies across tissues, diseases, or environmental contexts (Mancuso et al., 2019; Bryois et al., 2020).

To address these limitations, we propose Aggregated-TWAS (A-TWAS), an omnibus framework that integrates multiple Bayesian models which better capture the diverse and complex relationship between genotype and gene expression through two main enhancements. For the first enhancement, inspired by the flexibility of neuronized priors in Bayesian regression (Shin and Liu, 2021), A-TWAS aggregates information from multiple imputation models, each assuming different underlying structures for the cis-eQTL effect sizes. Specifically, we incorporate Bayesian Lasso [20], Horseshoe [7], and Horseshoe+ priors [4] to represent varying degrees of sparsity and shrinkage in the cis-eQTL effect size distribution. These priors are particularly well-suited for high-dimensional transcriptomic data, as they assume that only a subset of genetic variants have strong regulatory effects on gene expression, aligning with the sparse nature of transcriptomic regulation.

The second enhancement is that we aggregate the association p-values obtained from multiple imputation models using the Adaptive Combination of Evidence Test (ACAT) [15] in the second stage of A-TWAS, producing a unified and comprehensive gene-trait association p-value. This approach combines the strengths of different Bayesian models while mitigating the biases introduced by any single model’s assumptions. By leveraging the complementary strengths of multiple priors and integrating their results in a statistically principled manner, A-TWAS provides a more robust and flexible framework for TWAS analysis, particularly when the genetic architecture of gene expression is complex or varies across tissues and traits.

To evaluate the performance of A-TWAS, we conducted extensive simulations and real-world data analyses. The simulations demonstrate that A-TWAS improves imputation accuracy and statistical power while maintaining type-I error at a controlled level. Using GWAS summary statistics for schizophrenia and obsessive-compulsive disorder (OCD), we show that A-TWAS identifies substantially more disease-relevant genes compared to single-model TWAS approaches. Our results highlight the adaptability and robustness of A-TWAS in capturing diverse genetic architectures and underscore its utility as a powerful tool for gene-trait association studies.

### 2 Methods and Material

### 2.1 Overview of TWAS

Traditional TWAS methods mainly consist of the imputation step and the association step. The first imputation step utilizes individual-level genotype and transcriptome data from a reference panel to fit a regression model for each gene *g*, estimating the effect size of gene *g*’s cis-SNP on the expression level. Concretely, for each gene *g*, consider the following model:

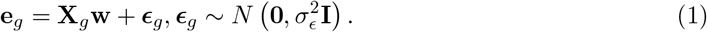

where **e**_*g*_ is an *n*-vector denoting the expression levels for gene *g*. **X**_*g*_ is an *n* × *m* genotype matrix recording the information of *m* cis-SNPs for gene *g*, only cis-SNPs within 1 MB of the flanking 5’ and 3’ ends are included. **w** is the effect size vector for *m* cis-SNPs of gene *g*, ***ϵ***_*g*_ is the random error term following a normal distribution. The intercept term is ignored in the model since **e**_*g*_ and **X**_*g*_ are assumed to be standardized.

In the second step of TWAS, if individual-level GWAS data is available, by using the effect size 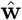 estimated from the first step, the genetically regulated expression (GReX) for an independent GWAS data can be imputed by 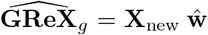, where **X**_new_ is the genotype information for gene *g* in the GWAS data. Then a generalized linear model (GLM) can be adopted to model the association between 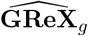 and phenotype **Y**:

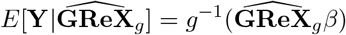

where *g* is the link function. For a quantitative phenotype **Y**, *g* can be the identity function; for a dichotomous phenotype *Y* , *g* can be set as the logit function. The association step evaluates the gene-level association by testing *H*_0_ : *β* = 0 versus *H*_1_ : *β* ≠ 0. If only summary-level GWAS data is available, the S-PrediXcan [1] test statistics could be used:

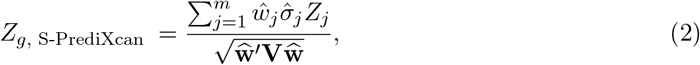

where *Z*_*j*_ is the *Z*-score test statistic for the *j*-th cis-SNP of gene *g* from summary-level GWAS data, **w** is estimated from the TWAS first step, *σ*_*j*_ is the standard deviation of the genotype information for the *j*-th cis-SNP, **V** is the genotype covariance matrix of gene *g*’s cis-SNPs, which could be approximated from reference panel used in the imputation step or other reference database from the same ancestry.

### 2.2 Baseline method

In stage I TWAS, multiple methods can be used to estimate the cis-eQTL effect size. In PrediXcan [10], Elastic-net [26] is adopted for imputation. Elastic-net is a regularized regression method that addresses sparsity by adding *L*_1_ and *L*_2_ norms as penalties. The Elastic-net model is defined as follows:

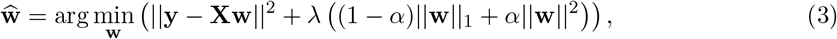

where *α* is a hyperparameter with value lies in [0, 1] that adjusts the shape of the penalty, and λ is a parameter controlling the strength of the penalty. In this work, *α* is manually set to 0.5, and λ is determined through cross-validation. Based on using Elastic-net for TWAS stage I imputation, the method employing GWAS summary data for stage II association testing via Equation 2 is S-PrediXcan [1], which is chosen as the baseline method in our study.

### 2.3 Bayesian Lasso

Unlike Elastic-net, which originates from the frequentist perspective, Bayesian Lasso [20] is a penalization method derived from the Bayesian perspective. It introduces a Laplace prior distribution on the weights for inference. In the context of this research, Bayesian Lasso places a Laplace prior on the cis-eQTL weights:

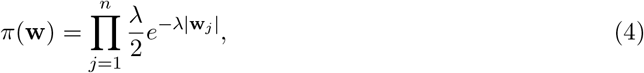

where λ is the hyperparameter controlling the strength of shrinkage. This parameter can be estimated using a Gibbs sampler and does not need to be set manually.

### 2.4 Horseshoe Method

The horseshoe prior developed by [7] is another powerful tool in high-dimensional regression and variable selection tasks. It exhibits strong shrinkage for small or irrelevant signals, effectively pulling them toward zero, while preserving large signals by means of the heavy tails. Moreover, unlike traditional Bayesian variable selection methods that typically impose discrete priors, which can be computationally expensive and challenging to apply in high-dimensional settings, the horseshoe prior sidesteps this issue by treating sparsity as a continuous property, enabling seamless inference without requiring hard thresholding. Specifically, the horseshoe model in the TWAS imputation step can be formulated as:

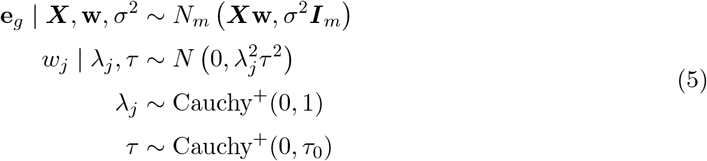

where Cauchy^+^(0, *a*) is the standard half-Cauchy distribution defined on *R*^+^ with scale parameter *a*. λ_*j*_ is the local shrinkage parameter and *τ*_0_ is the global shrinkage parameter.

### 2.5 Horseshoe+ Method

The Horseshoe+ prior[4] was proposed based on Horseshoe. It extends the horseshoe prior by introducing an additional layer of hierarchy to further adapt the shrinkage behavior. Assuming the model as 1 after standardization for each term, the horseshoe+ prior for the cis-eQTL weights is:

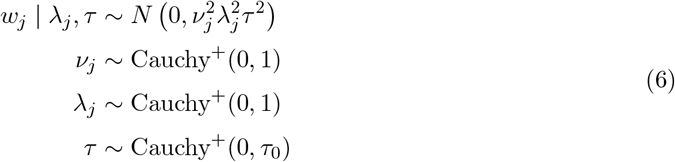

here *ν*_*j*_ is a new auxiliary shrinkage parameter, allowing finer control of shrinkage at the local level, offering greater adaptability for different sparsity patterns. Since the SNP signals are usually very sparse in the genes, Horseshoe and Horseshoe+ are naturally a good choice for imputing the cis-eQTL weights in the first stage of TWAS. The solution of Horseshoe and Horseshoe+ models could be attained by reformulating them into hierarchical models, and calculating the posterior estimation through Monte Carlo Markov Chain(MCMC).

### 2.6 A-TWAS and ACAT Test

We introduce Aggregated-TWAS (A-TWAS), a framework that integrates multiple Bayesian models to more effectively capture the complex and diverse relationships between genotype and gene expression. A-TWAS first involves training imputation models using Bayesian Lasso, Horseshoe, and Horseshoe+ for each gene. Next, the association between genotype and phenotype is tested using Equation 2 for each set of cis-eQTL weights, resulting in a set of TWAS *p*-values corresponding to each Bayesian method.

To integrate the *p*-values obtained from the three Bayesian imputation methods, we employ ACAT [15, 16]. Let *p*_*i*_ represent the p-value from the *i*-th Bayesian method, and let *d* denote the total number of Bayesian methods used for integration. The Cauchy combination test statistics of ACAT is defined as:

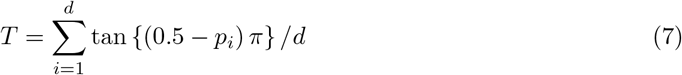

Since the null distribution of *T* can be well approximated by a Cauchy distribution under arbitrary dependency structures for *p*_*i*_, the combined *p*-value for three Bayesian methods can be calculated as 0.5 − arctan(*T* )*/π*. This combined *p*-value returned by the ACAT test is just the TWAS *p*-value for our A-TWAS method.

### 2.7 ROS/MAP Dataset

For the real data analysis, we performed the first imputation step of TWAS on the Religious Orders Study and Memory and Aging Project (ROS/MAP) dataset[3][2]. We used the R package “bigsnpr” to preprocess the genotype data, retaining variants with a minor allele frequency (MAF) > 5% and a Hardy-Weinberg equilibrium Fisher’s exact test *p*-value < 1 × 10^−7^. With respect to the gene expression data, for a specific gene, if more than 70% of the samples in the cohort had low FPKM values (the threshold is set to 1 in this study) for a given gene, we excluded that gene from the TWAS analysis. Furthermore, we used linear regression to adjust for confounding factors, including sex, age, study cohort (ROS or MAP), and the top three principal components of the genotype data. After data preprocessing, a total of 10,940 genes were included for the imputation step of the TWAS analysis.

## 3 Simulation Study

To evaluate the performance of Bayesian Lasso, Horseshoe and Horseshoe+, together with their aggregated performance(A-TWAS), we conducted simulation experiments to compare their test *R*^2^, TWAS power, and type-I error. We utilized genotype information from the ROS/MAP dataset by using a randomly selected gene *CCDC*104 from chromosome 2 to construct the genotype matrix in simulation, while the gene expression and phenotype data were artificially simulated. 50% of the samples were randomly selected for training, and the remaining 50% were used for evaluating the imputation performance by test *R*^2^.

We varied the fraction of causal SNPs *p*_*cs*_ (0.01, 0.05, 0.1, 0.3) and gene expression heritability 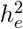 (0.01, 0.05, 0.1) to simulate gene expression data, where the gene expression heritability refers to the proportion of variance explained by causal SNPs. Under the model 1, 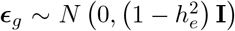 , the effect size for causal SNPs are sampled from *N* (0, 1), and then we re-scaled the effect size to ensure the expression heritability to be 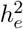.

In order to access the performance of TWAS power, we assumed the phenotype data are simulated by

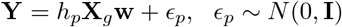

where 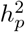 is the phenotypic heritability, namely the proportion of phenotypic variance explained by gene expression. We set 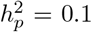 in this study. Under this model assumption for the phenotype data, GWAS *z*-scores of the summary statistics used in stage II of TWAS are generated from a multivariate normal distribution 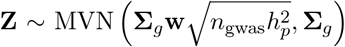 [8], whereΣ_*g*_ is the correlation matrix of the standardized genotype matrix, and *n*_gwas_ is the assumed GWAS sample size, which is varied with value *n*_gwas_ = (20000, 50000, 80000) in this study. Each simulation scenario was repeated 500 times.

To evaluate the type I error of each method, we assumed the phenotype was independent of gene expression and generated the phenotype directly from *N* (0, 1). For each 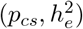 scenario, the phenotype was simulated with 500 replications.

### 3.1 Simulation Result

For the simulation results, A-TWAS achieves the highest statistical power under most scenarios when the significance threshold for the power calculation is *p* = 0.05*/*500 = 1×10^−4^ (adjusted using the Bonferroni correction). Details of the simulation results can be found in Supplementary Figs. S1, S2, S3, and S4. A-TWAS robustly outperforms the baseline S-PrediXcan method and is slightly inferior to other individual Bayesian methods only in a few cases, highlighting the effectiveness of A-TWAS.

For the individual Bayesian methods, they consistently achieve higher power than S-PrediXcan. When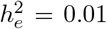, Horseshoe and Horseshoe+ outperform Bayesian Lasso, but their performance achieve a similar level when 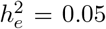 and 0.1. All methods exhibit comparable performance in test *R*^2^. Only when *p*_*cs*_ ≥ 0.05 and 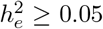, the Bayesian methods perform slightly better than S-PrediXcan, which employs the frequentist Elastic-net.

Regarding type-I error, the error rate is defined as the proportion of TWAS p-values falling below the threshold *p* = 0.05. Among the methods, the type-I error rates for all Bayesian methods are similar, while the rate for S-PrediXcan is stable and slightly lower than that of the Bayesian methods. Furthermore, the aggregation of Bayesian methods (A-TWAS) increases power without inflating the type-I error. Overall, all methods maintain an acceptable type-I error rate.

## 4 Real Application

To evaluate the performance of A-TWAS and the individual Bayesian methods on real data, we applied these methods, along with S-PrediXcan as a baseline, to identify significant genes. The ROS/MAP dataset was used to train GReX imputation models, and GWAS summary data for Schizophrenia [19] and Obsessive-Compulsive Symptoms [21] from the Psychiatric Genomics Consortium (PGC) were utilized for 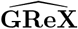 and phenotype association testing.

In stage I of TWAS, the genotype data used to train the GReX imputation model was derived from the intersection of SNPs in both the ROS/MAP dataset (quality-controlled as described in Section 2.7) and the GWAS summary statistics for the disease. The gene expression data also came from the ROS/MAP dataset with appropriate adjustments (detailed in Section 2.7). Half of the samples were used for training, and the other half were reserved for testing.

In stage II of TWAS, the genotype information for the entire sample set was used as the reference LD panel. Imputation models with test *R*^2^ > 0.01 were deemed effective and selected for the subsequent stage of TWAS. If the imputation model was null, both its train *R*^2^ and test *R*^2^ were set to 0.

### 4.1 Schizophrenia

The Schizophrenia GWAS summary dataset [19] contains 11,260 cases and 24,542 controls from the UK. We calculated its heritability and the proportion of causal SNPs using the two-component model of GENESIS [25] and found that the heritability (*h*^2^) and *p*_*cs*_ of this dataset are 0.707 and 0.0142, respectively. The number of genes containing SNP information in the Schizophrenia GWAS summary data is 10,893. Among these genes, 7,411 imputation models qualified for TWAS analysis using A-TWAS, and more than 6,000 models are qualified for Bayesian methods, compared to 5,341 models in S-PrediXcan (Table 1).

**Table 1:**
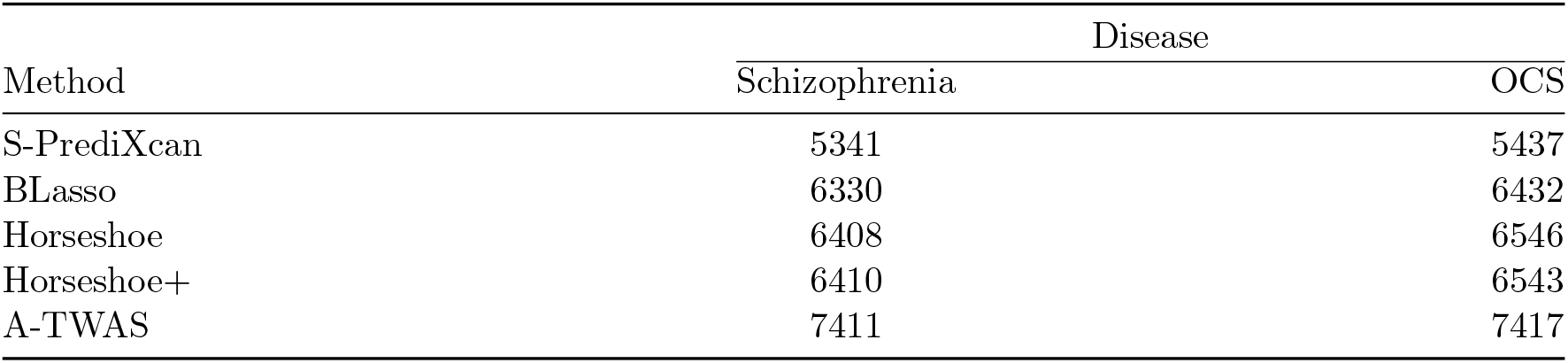
Effective Imputation Model Counts (test *R*^2^ > 0.01).

| Method | Disease |  |
| --- | --- | --- |
|  | Schizophrenia | OCS |
| S-PrediXcan | 5341 | 5437 |
| BLasso | 6330 | 6432 |
| Horseshoe | 6408 | 6546 |
| Horseshoe+ | 6410 | 6543 |
| A-TWAS | 7411 | 7417 |

S1, S2, S3, and S4. A-TWAS robustly outperforms the baseline S-PrediXcan method and is slightly inferior to other individual Bayesian methods only in a few cases, highlighting the effectiveness of A-TWAS.

Figure S5 shows the train *R*^2^ and test *R*^2^ results for Schizophrenia. The train *R*^2^ values for Bayesian methods are significantly higher than those for S-PrediXcan. For test *R*^2^, all methods perform similarly to each other, with Horseshoe and Horseshoe+ slightly outperforming the others, which aligns with our simulation results.

In the TWAS results, A-TWAS identified 89 significant genes with p-values reaching the Bonferroni-corrected significance level, the highest among all methods. Given the 10,893 genes containing SNP information in the Schizophrenia dataset, the corrected p-value threshold is 4.59 × 10^−6^. The number of significant genes identified by other methods is as follows: S-PrediXcan 75, BLasso 43, Horseshoe 79, and Horseshoe+ 77. The Manhattan plot for A-TWAS results is shown in Fig. 1, while Manhattan plots for other methods are provided in Supplementary Fig. S6.

**Figure 1:**
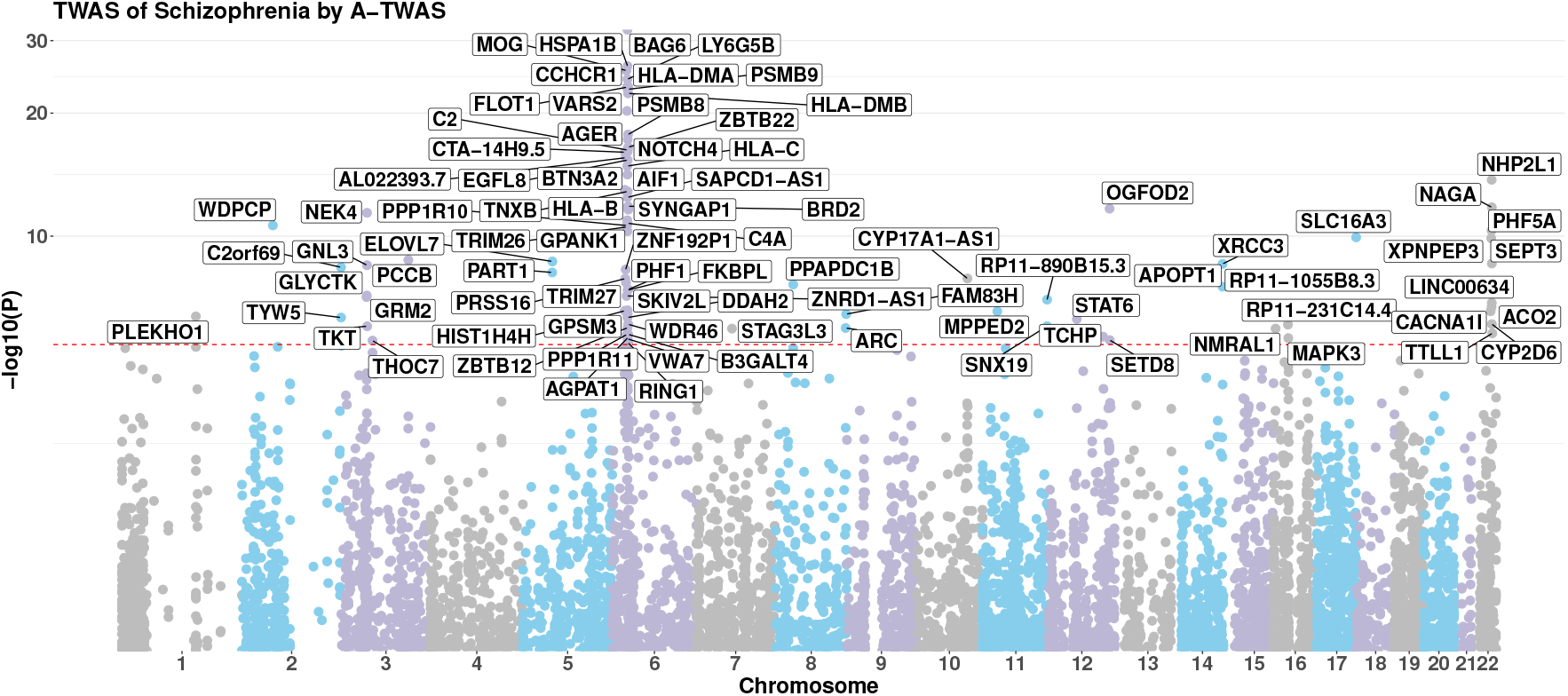
Manhattan plot for Schizophrenia.

Among the genes identified by A-TWAS, 24 are listed in the GWAS Catalog under the trait “Schizophrenia”, the most among all methods, followed by Horseshoe method (S-PrediXcan 23, BLasso 12, Horseshoe 20, Horseshoe+ 18). Details of these results are presented in Supplementary Table S1. Enrichment analysis of the significant genes identified by A-TWAS successfully identified the Schizophrenia pathway (KEGG disease) with a p-value of 0.0295, further demonstrating the efficacy of A-TWAS.

### 4.2 Obsessive-Compulsive Symptoms

The study of Obsessive-Compulsive Symptoms (OCS) includes 33,943 samples of European ancestry [21]. The GWAS results from this study found no SNPs or genes significantly associated with OCS, as none reached the Bonferroni-corrected significance level. Using GENESIS, the *p*_*cs*_ and heritability for this GWAS summary data were calculated to be 8.592×10^−4^ and 0.0133, respectively. There are 11,014 genes containing SNP information in the OCS dataset. Among these genes, 5,437 attained valid imputation models using S-PrediXcan, which is fewer than the 7,417 identified by A-TWAS and >6,000 models identified by Bayesian methods (Table 1).

The patterns of train *R*^2^ and test *R*^2^ (Supplementary Fig. S8) for the OCS data are similar to those observed for Schizophrenia, as the training sources for the imputation models are largely the same. The only difference arises from variations in the SNPs recorded in the two sets of summary statistics.

In the TWAS results, A-TWAS successfully identified five new significant genes for OCS (Fig. 2), while S-PrediXcan identified none. The corrected p-value threshold is 4.54×10^−6^. Manhattan plots for the other methods are shown in Supplementary Fig. S9, and the p-values for the significant genes are provided in Supplementary Table S2.Overall, these results from real data demonstrate the effectiveness of A-TWAS.

**Figure 2:**
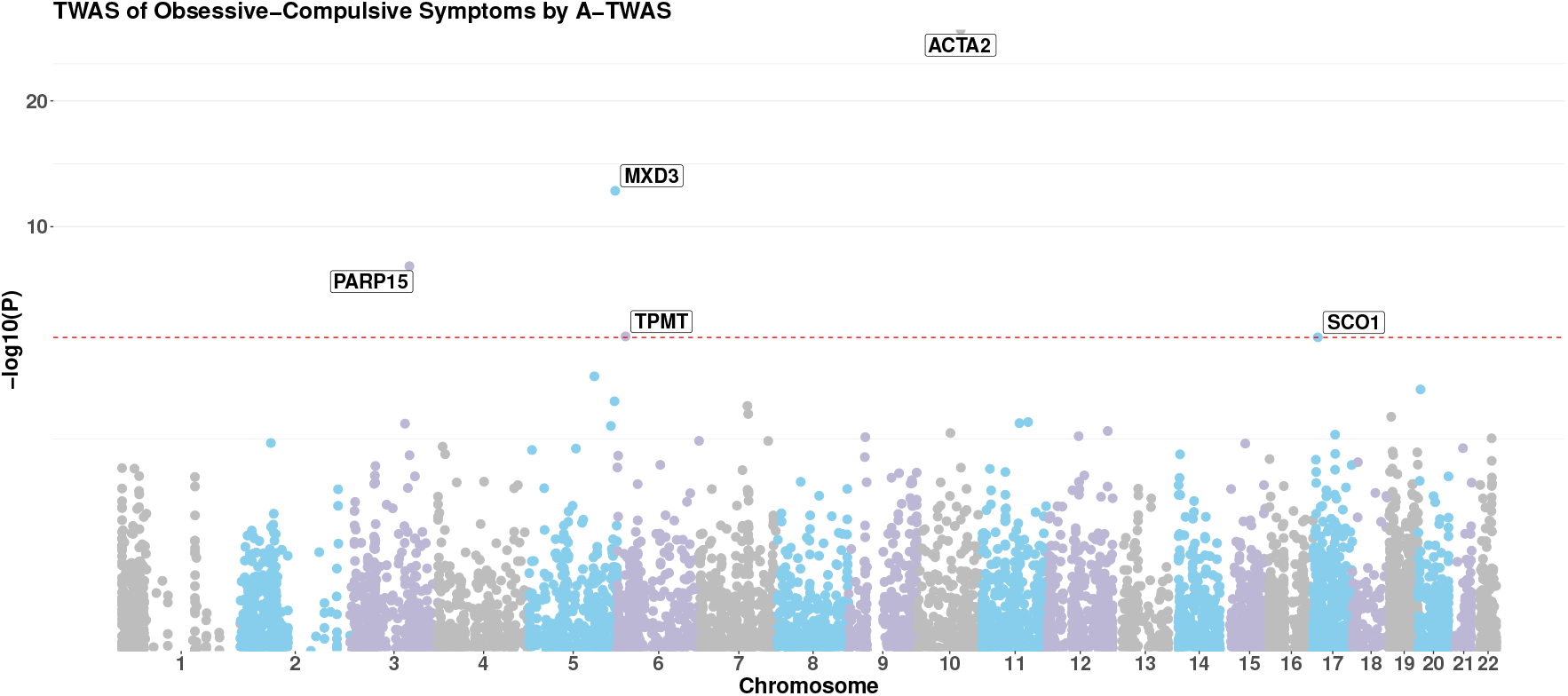
Manhattan plot for Obsessive-Compulsive Symptoms.

## 5 Conclusion and Discussion

In this work, A-TWAS was proposed to better capture the complex relationship between genotype and gene expression by combining multiple imputation models with different Bayesian priors. It enhances the detection of genetic risk factors by integrating various Bayesian models, improving power over the baseline method. We chose Bayesian Lasso, Horseshoe, and Horseshoe+ as the individual imputation models for combination after proving their efficacy. Detailed simulation experiments and real data analysis were performed for A-TWAS, together with each single Bayesian method and the baseline method S-PrediXcan, to investigate their performance.

In the simulation study, A-TWAS robustly outperformed other methods regardless of the underlying genetic architecture of the data, demonstrating the advantage of the omnibus method in capturing a more comprehensive genetic structure compared to single methods. For the single methods, Horseshoe and Horseshoe+ performed better than Bayesian Lasso when the causal SNPs in the gene were sparse. All these methods outperform the baseline S-PrediXcan. For test *R*^2^, no significant differences were observed between the methods. Regarding type-I error, although A-TWAS and all Bayesian methods did not reach the performance of S-PrediXcan, when considering the balance between power and type-I error, all of these methods achieved relatively well-controlled type-I error rates.

For the real data analysis, GWAS summary data for the diseases Schizophrenia and Obsessive-Compulsive Symptoms (OCS) were adopted to test the performance of the methods. Among the methods, A-TWAS showed its advantage in that it could attain most of the imputation models enabling the next stage of TWAS, since it is the union of results attained by the three Bayesian methods. In Schizophrenia, A-TWAS successfully identified most of the significant genes; among the significant genes, the set of gene detected by A-TWAS includes the most that have been proven to be related to Schizophrenia and reported in the GWAS Catalog. The significant genes selected by A-TWAS also successfully identified the Schizophrenia disease pathway through enrichment analysis. For single Bayesian methods, although slightly inferior to A-TWAS, Horseshoe and Horseshoe+ have a comparable performance which is better than other single methods regarding these criteria. In the analysis of OCS, this group of GWAS summary data originally reported no significant SNPs or genes. By incorporating this data into TWAS methods, no significant genes were detected by S-PrediXcan and Bayesian Lasso. However, A-TWAS successfully identified 5 genes (Horseshoe 5 and Horseshoe+ 2). All of these evidences prove the superiority of using the continuous shrinkage prior Horseshoe and Horseshoe+ for imputation model, and the efficacy of the omnibus method A-TWAS.

From the results obtained in our work, A-TWAS clearly demonstrates its ability in identifying genetrait relationships. However, since all Bayesian methods obtain their solutions using MCMC, which requires a large number of iterations, the computation speed is much slower than the frequentist S-PrediXcan, limiting the efficiency of A-TWAS. For the implementation of A-TWAS (GitHub website: https://github.com/Yilan-Liang/A-TWAS.), our code provides functions for calculating imputation based on an individual Bayesian method or any combination of the Bayesian methods, enabling the trade-off between power and efficiency. Either the Horseshoe or Horseshoe+ model is recommended if only an individual model is required, as they perform robustly better than Bayesian Lasso.

## 6 Code and Data Availability

### 6.1 Code Availability

A-TWAS: https://github.com/Yilan-Liang/A-TWAS.

S-PrediXcan: https://github.com/hakyimlab/MetaXcan.

bayesreg (the R package bayesreg support the calculation of Bayesian Lasso, Horseshoe and Horse-shoe+): https://cran.r-project.org/web/packages/bayesreg/index.html.

### 6.2 Data Availability

The data that support the findings of this study can be downloaded from the following sources:

All protected data of the GTEx project are available through the database of Genotypes and Phenotypes (dbGaP) (accession number phs000424.v8.p2) at https://www.ncbi.nlm.nih.gov/projects/gap/cgi-bin/study.cgi?study_id=phs000424.v8.p2.

ROSMAP: https://www.synapse.org/#!Synapse:syn3219045.

GWAS summary statistics for Schizophrenia: https://figshare.com/articles/dataset/scz2018clozuk/14681220.

GWAS summary statistics for Obsessive-Compulsive Symptoms: https://figshare.com/articles/dataset/ocs2024/26352157.

## 7 Author contributions statement

Y.D.Z. conceived and designed the model. Y.L and H.W. designed the algorithm, implemented the software, conducted analysis of simulation and real data, and wrote the original draft of the manuscript. Y.D.Z. supervised the study and contributed to the critical revision of the manuscript.

Y.L. and H.W. made equal contributions.

## 8 Acknowledgments

This work was supported, in part, by Hong Kong Research Grants Council General Research Fund [17307324]. The Genotype-Tissue Expression (GTEx) Project was supported by the Common Fund of the Office of the Director of the National Institutes of Health, and by NCI, NHGRI, NHLBI, NIDA, NIMH, and NINDS.

## A Supplemental Figures

**Figure S1:**
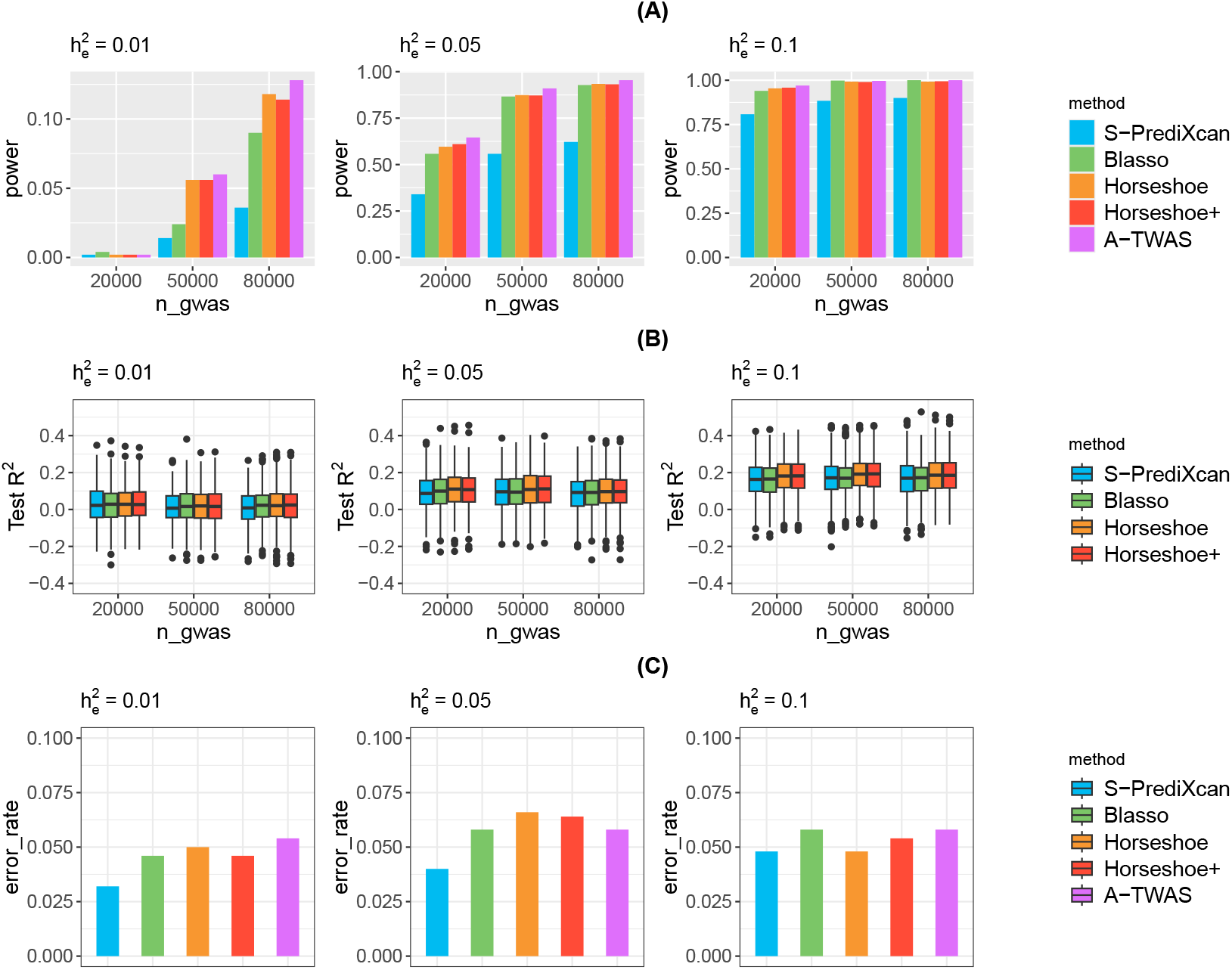
simulation result under p_cs_ = 0.01 scenario. **(A)** TWAS power with threshold of p-value < 10^−4^ (Bonferroni corrected significance level) vary by *h*_*e*_ and GWAS sample size given by S-PrediXcan and all proposed Bayesian method. **(B)** The test *R*^2^ of S-PrediXcan, Bayesian Lasso, Horseshoe and Horseshoe+ in stage I TWAS vary by *h*_*e*_ and GWAS sample size. **(C)** The box plot of the p-value under the simulation of type-one error evaluation.

**Figure S2:**
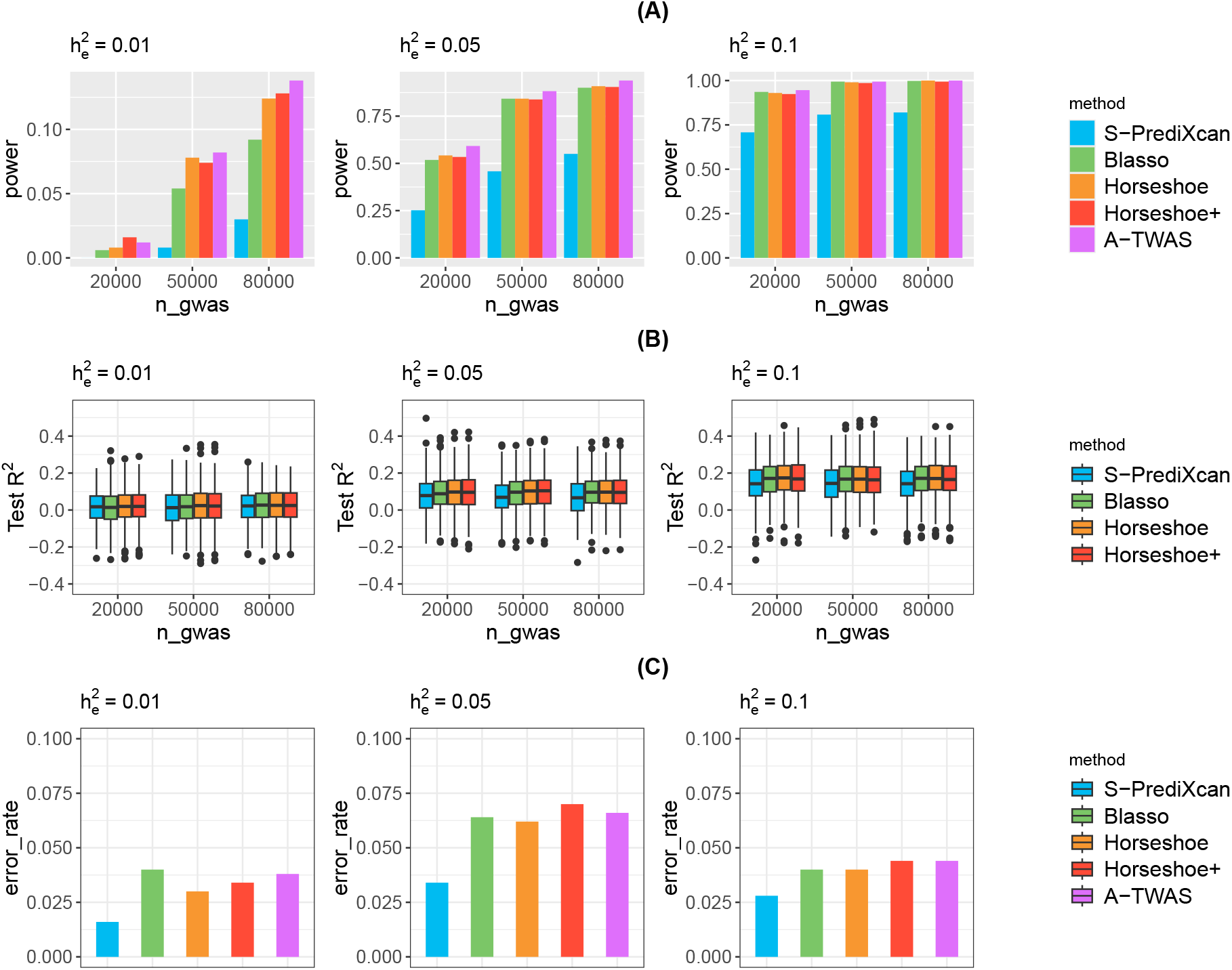
simulation result under p_cs_ = 0.05 scenario. **(A)** TWAS power with threshold of p-value < 10^−4^ (Bonferroni corrected significance level) vary by *h*_*e*_ and GWAS sample size given by S-PrediXcan and all proposed Bayesian method. **(B)** The test *R*^2^ of S-PrediXcan, Bayesian Lasso, Horseshoe and Horseshoe+ in stage I TWAS vary by *h*_*e*_ and GWAS sample size. **(C)** The box plot of the p-value under the simulation of type-one error evaluation.

**Figure S3:**
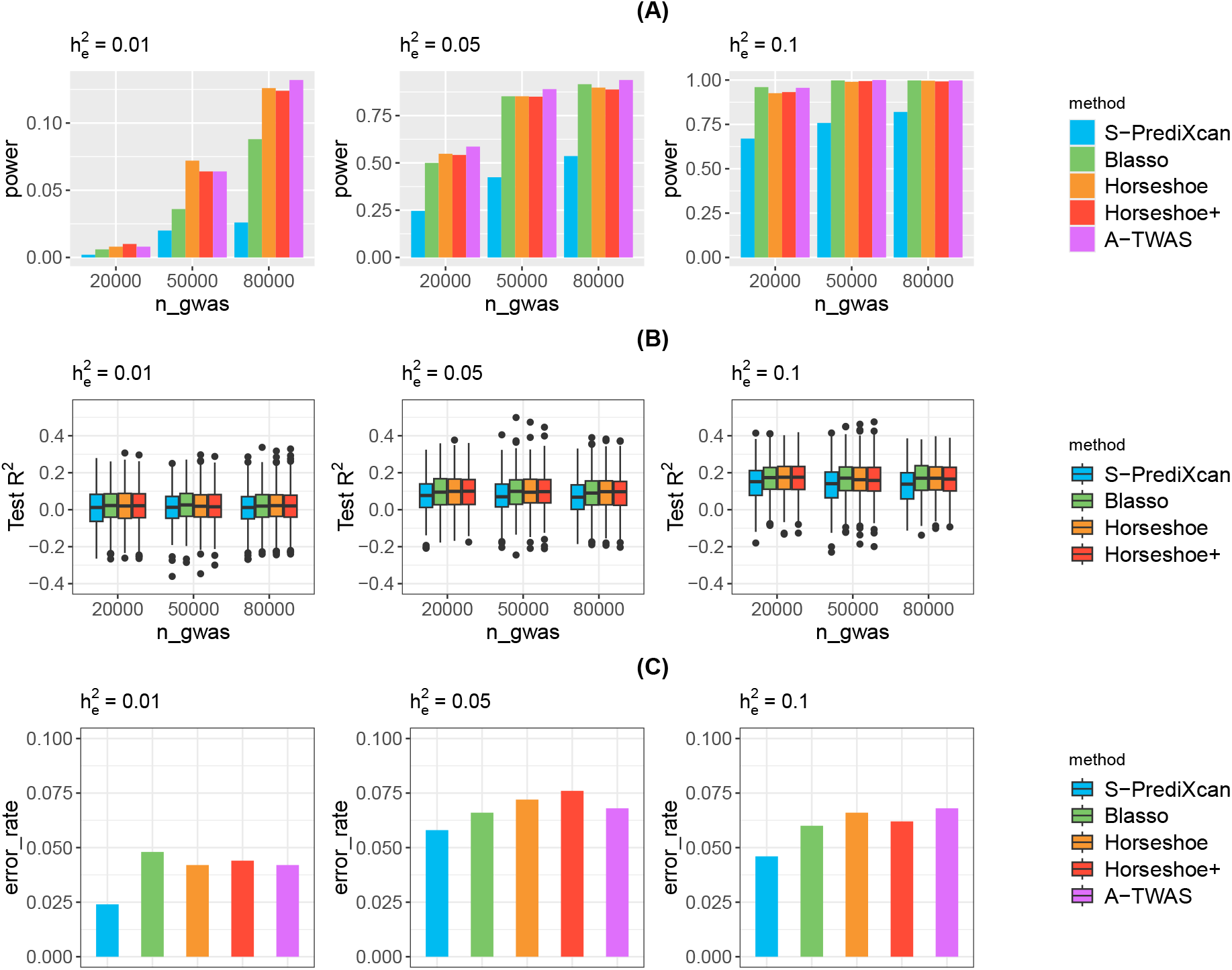
simulation result under p_cs_ = 0.1 scenario. **(A)** TWAS power with threshold of p-value < 10^−4^ (Bonferroni corrected significance level) vary by *h*_*e*_ and GWAS sample size given by S-PrediXcan and all proposed Bayesian method. **(B)** The test *R*^2^ of S-PrediXcan, Bayesian Lasso, Horseshoe and Horseshoe+ in stage I TWAS vary by *h*_*e*_ and GWAS sample size. **(C)** The box plot of the p-value under the simulation of type-one error evaluation.

**Figure S4:**
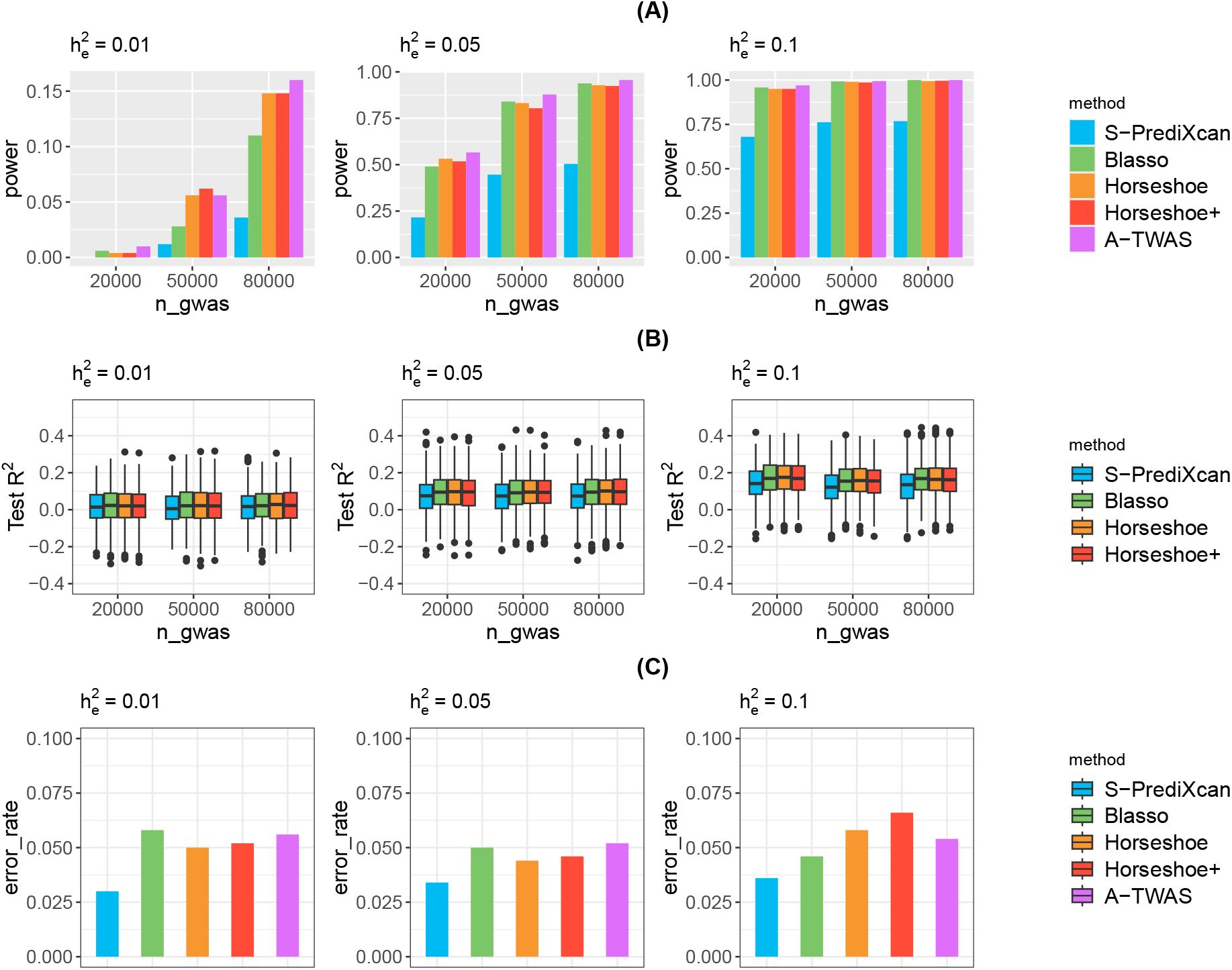
simulation result under p_cs_ = 0.3 scenario. **(A)** TWAS power with threshold of p-value < 10^−4^ (Bonferroni corrected significance level) vary by *h*_*e*_ and GWAS sample size given by S-PrediXcan and all proposed Bayesian method. **(B)** The test *R*^2^ of S-PrediXcan, Bayesian Lasso, Horseshoe and Horseshoe+ in stage I TWAS vary by *h*_*e*_ and GWAS sample size. **(C)** The box plot of the p-value under the simulation of type-one error evaluation.

**Figure S5:**
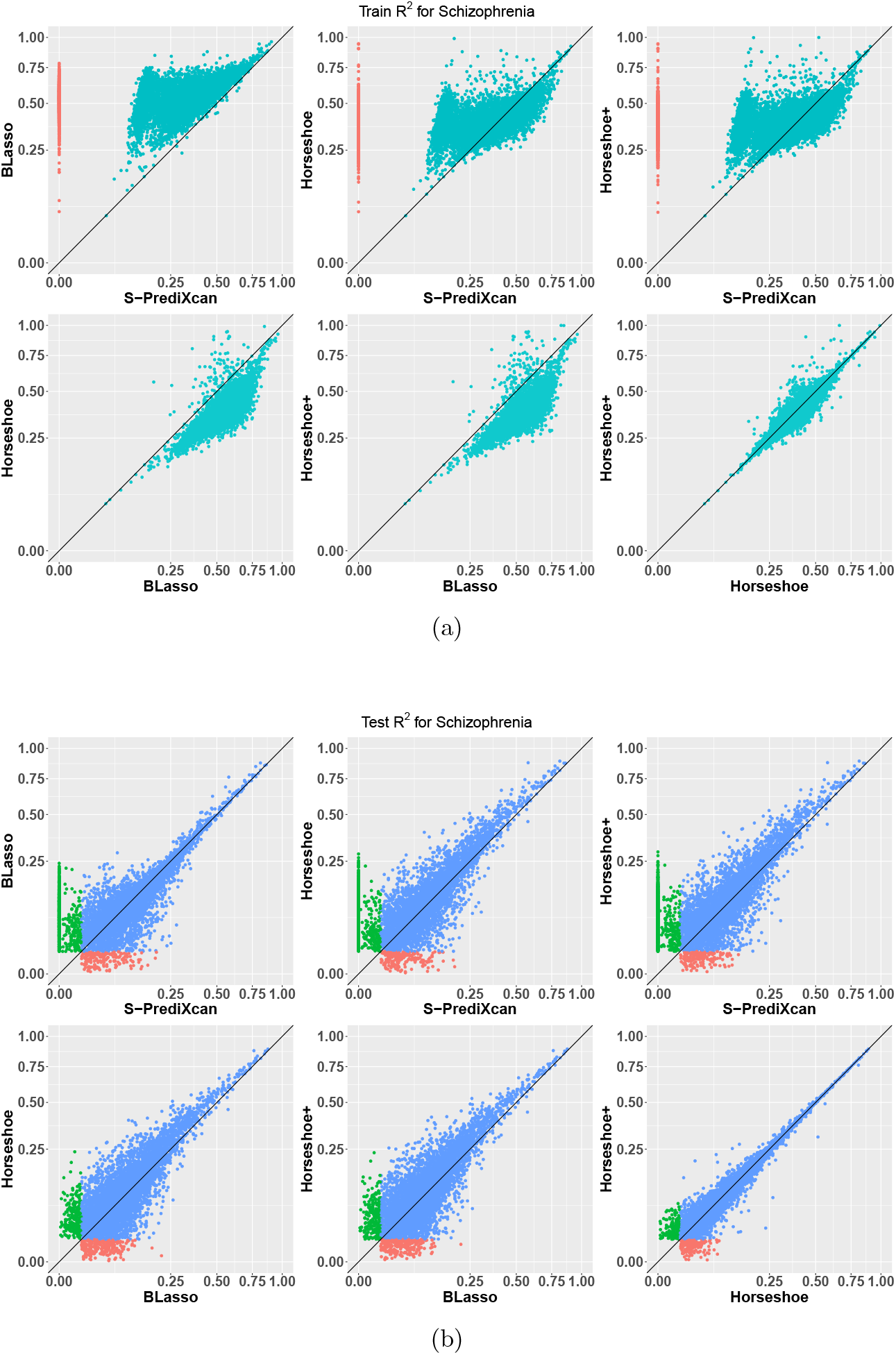
test R^2^ and train R^2^ for Schizophrenia. In train *R*^2^, the red points indicating models with *R*^2^ > 0.01 for method on y-axis while < 0.01 for method on x-axis; the blue points indicating models with *R*^2^ > 0.01 for methods on both axis. In test *R*^2^, the red points indicating models with *R*^2^ > 0.01 for method on x-axis while < 0.01 for method on y-axis; the green points indicating models with *R*^2^ > 0.01 for method on y-axis while < 0.01 for method on x-axis; and the green points indicating models with *R*^2^ > 0.01 for methods on both axis.

**Figure S6:**
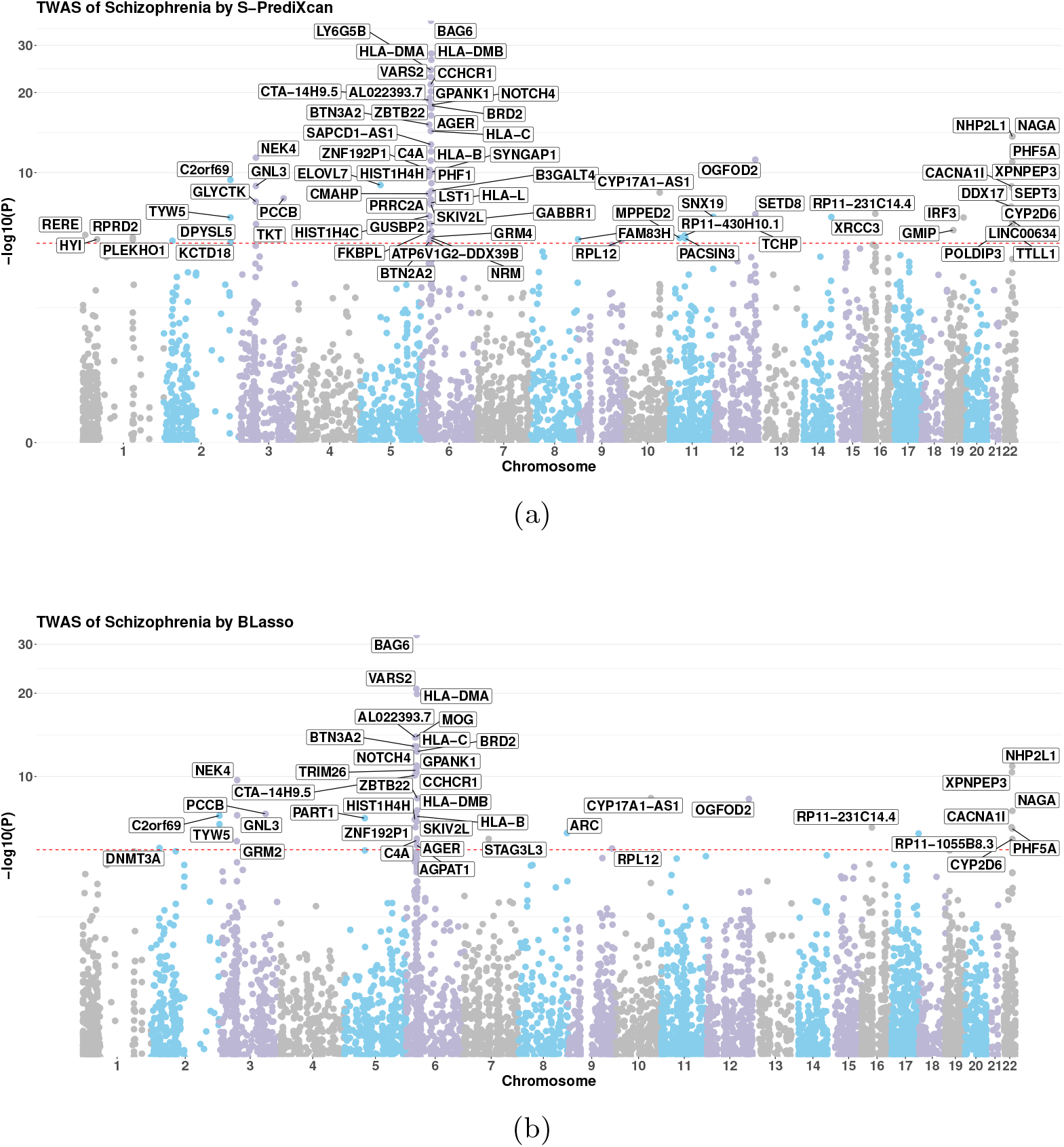
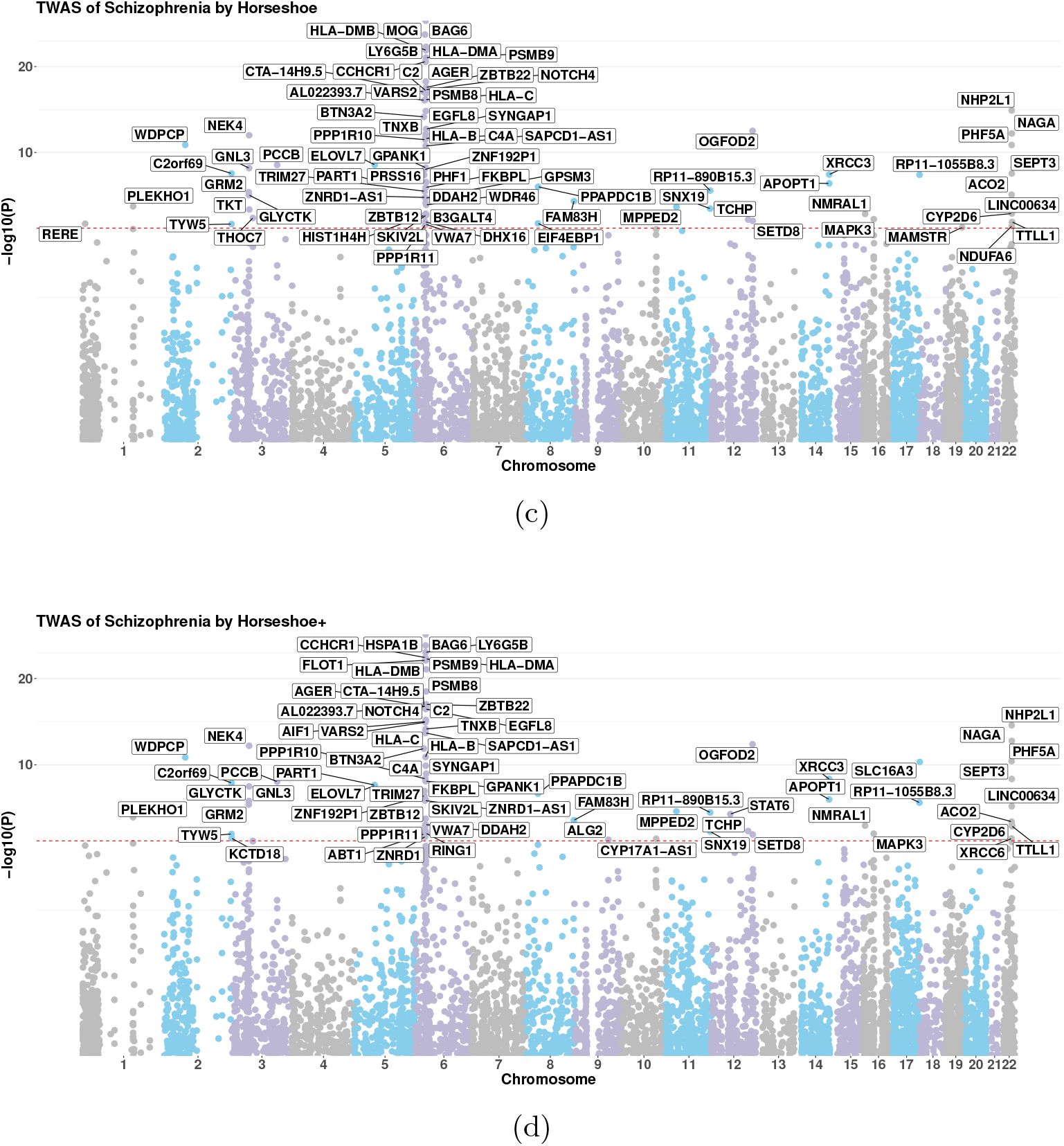
Manhattan plots for Schizophrenia.

**Figure S7:**
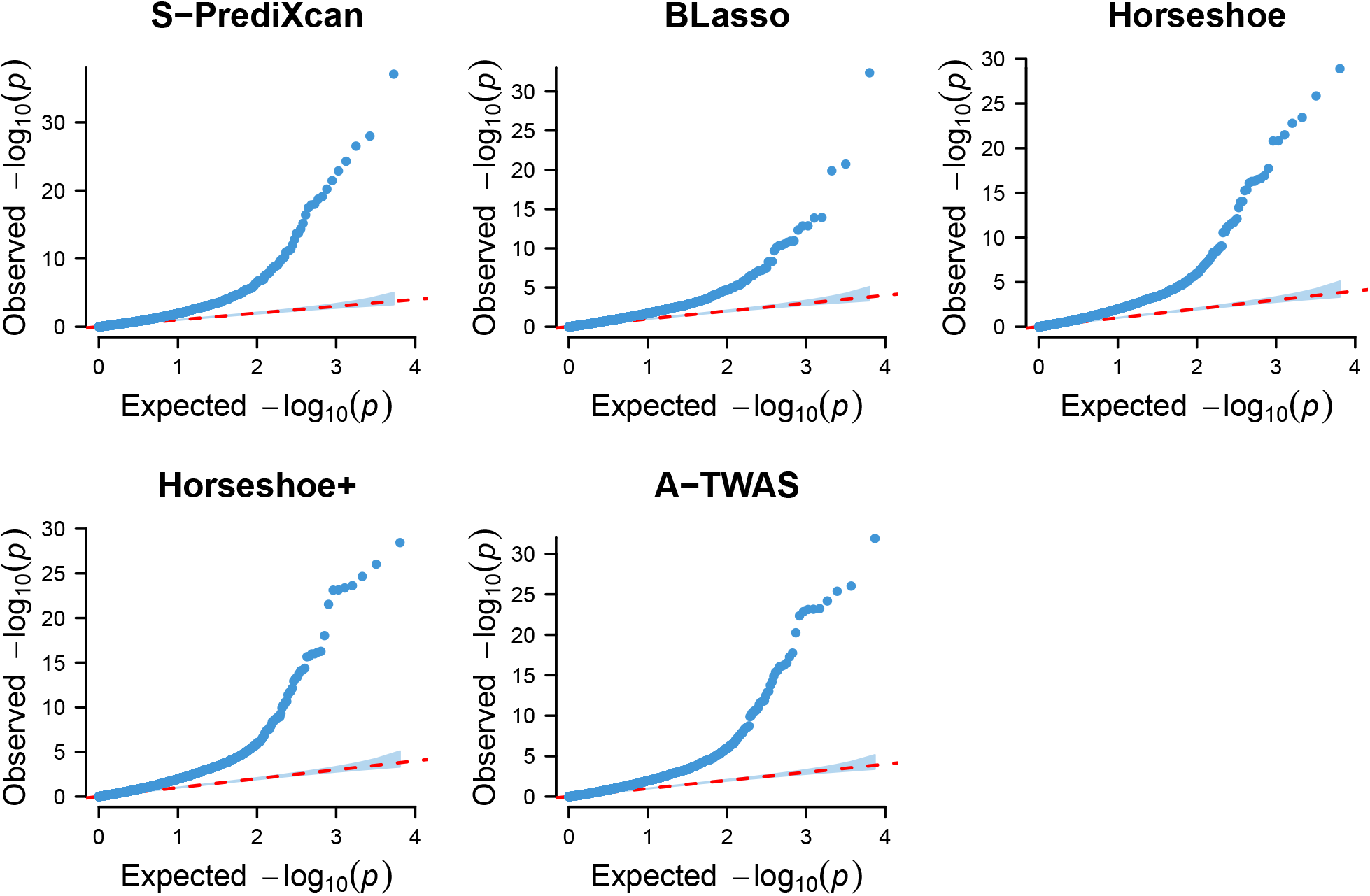
Quantile–Quantile plot of TWAS p–values under uniform distribution from Schizophrenia.

**Figure S8:**
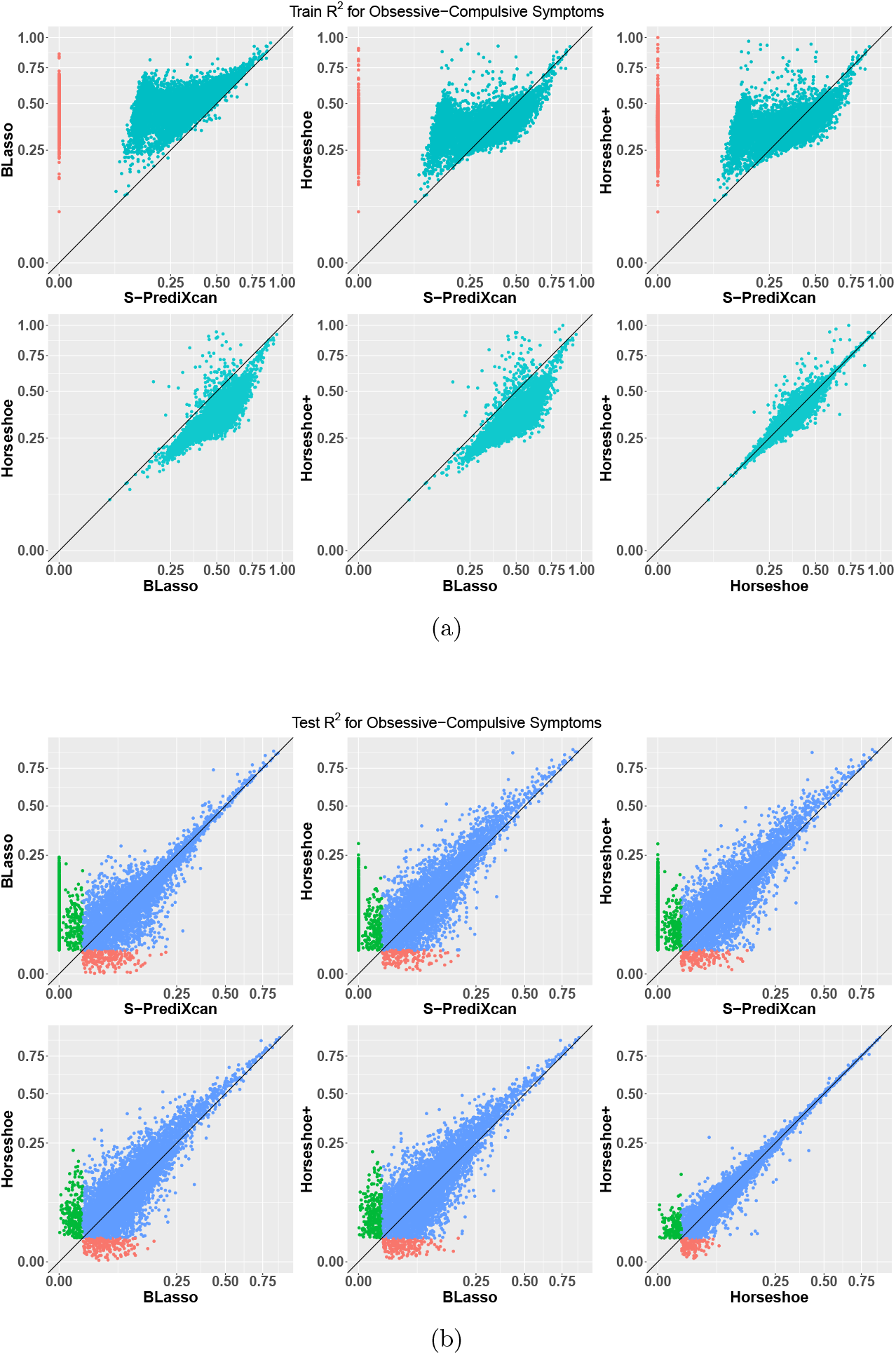
test R^2^ and train R^2^ for Obsessive-Compulsive Symptoms. In train *R*^2^, the red points indicating models with *R*^2^ > 0.01 for method on y-axis while < 0.01 for method on x-axis; the blue points indicating models with *R*^2^ > 0.01 for methods on both axis. In test *R*^2^, the red points indicating models with *R*^2^ > 0.01 for method on x-axis while < 0.01 for method on y-axis; the green points indicating models with *R*^2^ > 0.01 for method on y-axis while < 0.01 for method on x-axis; and the green points indicating models with *R*^2^ > 0.01 for methods on both axis.

**Figure S9:**
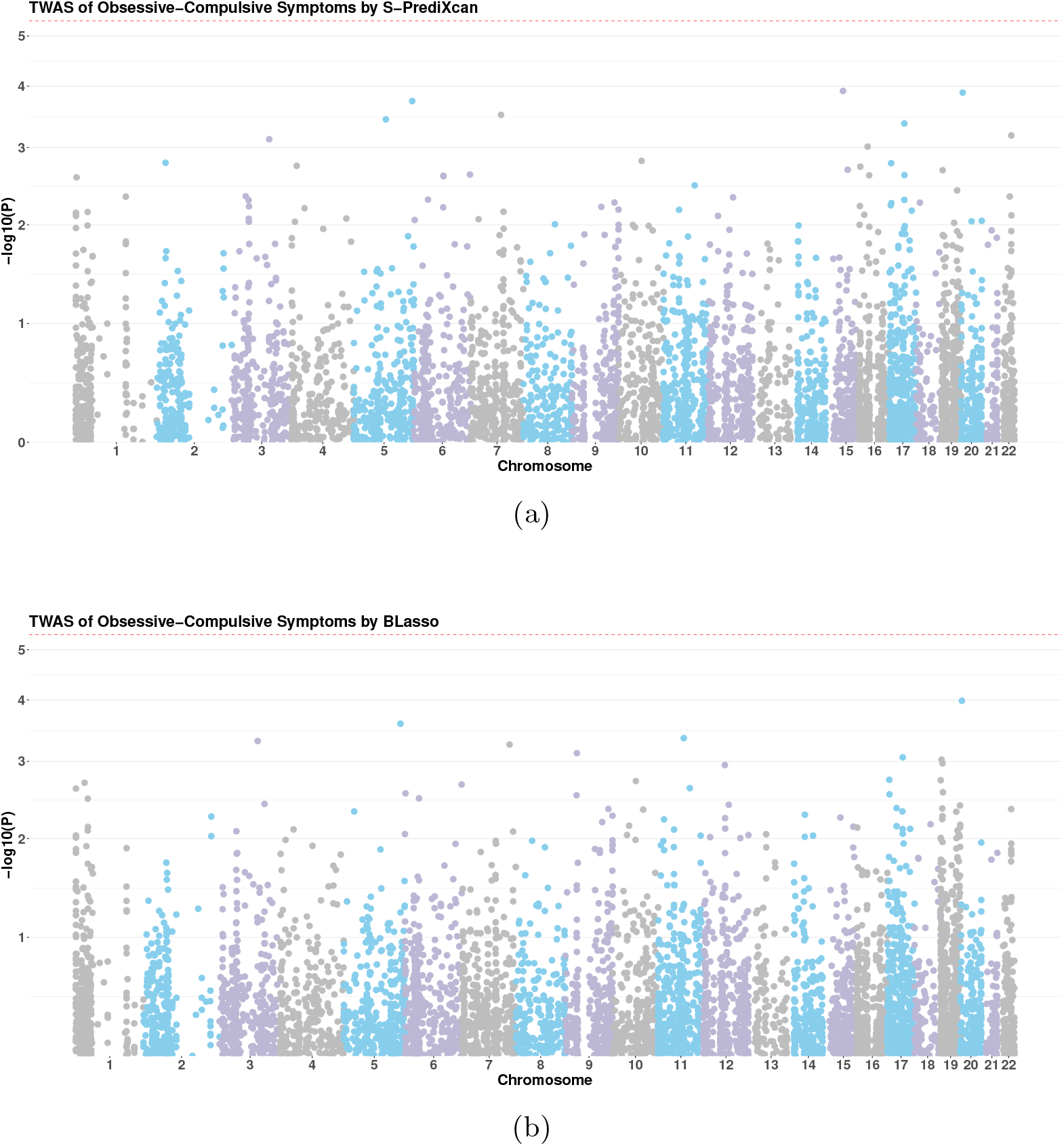
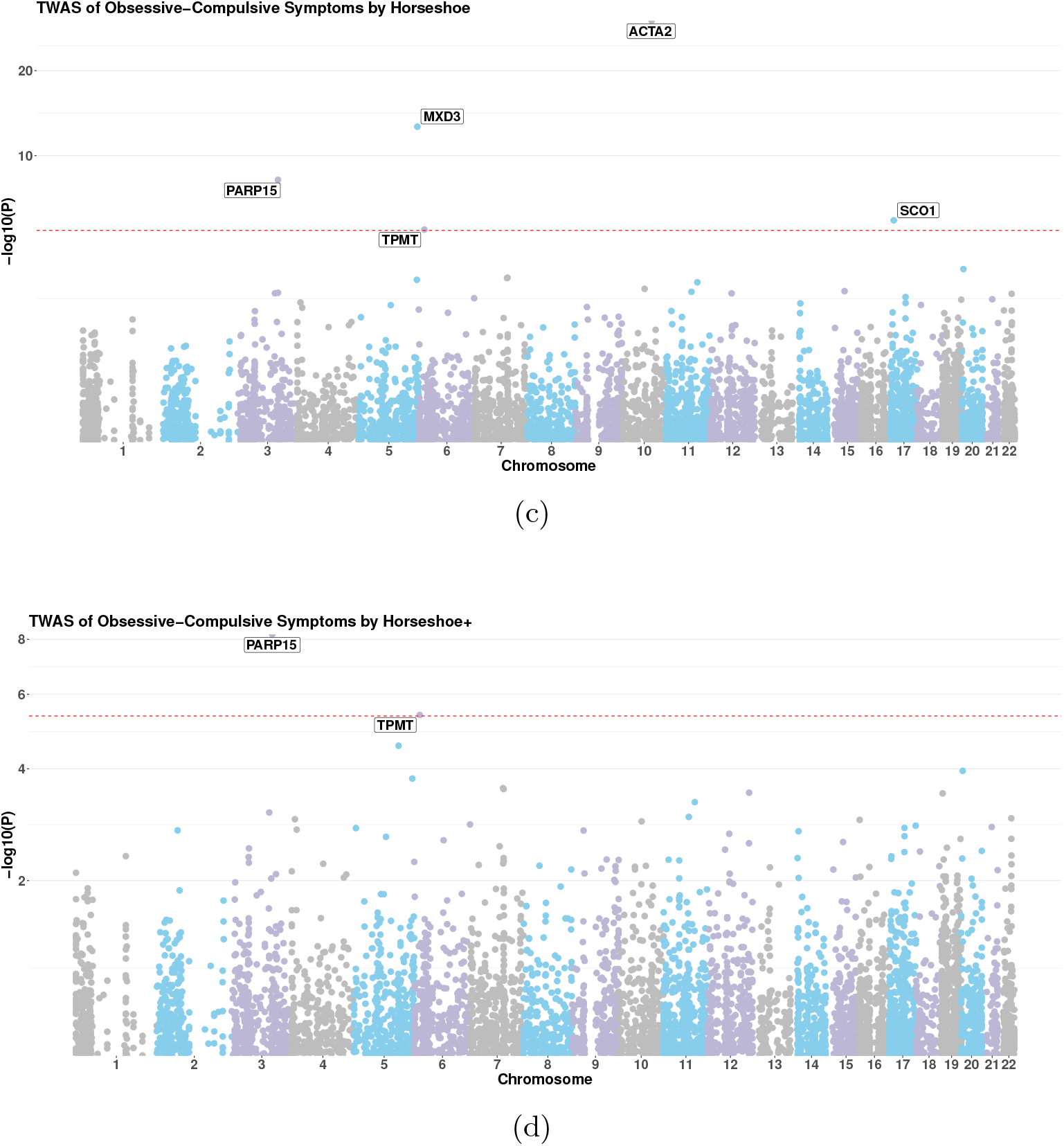
Manhattan plots for Obsessive–Compulsive Symptoms.

**Figure S10:**
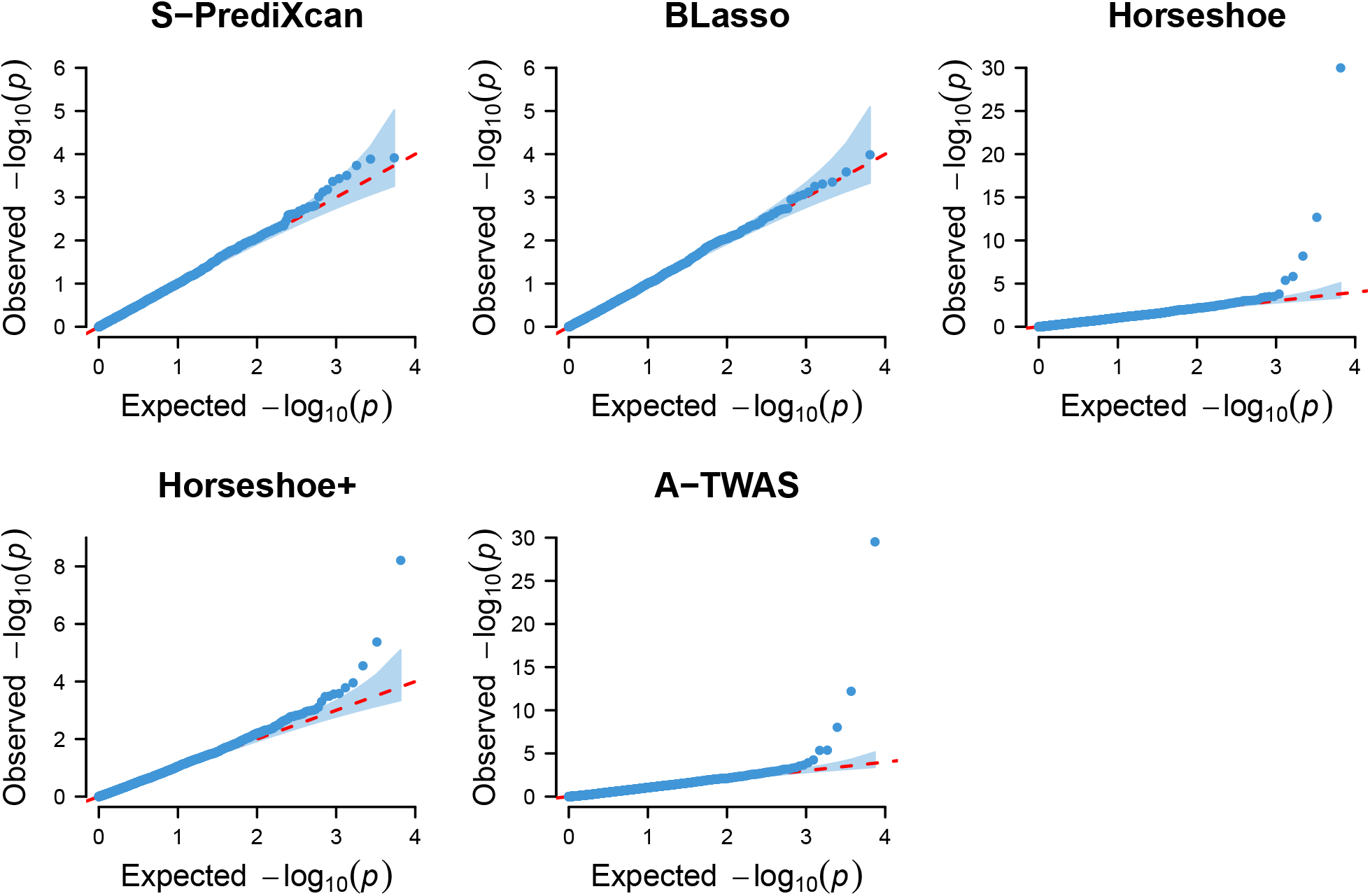
Quantile–Quantile plot of TWAS p–values under uniform distribution from Obsessive–Compulsive Symptoms.

## B Supplemental Tables

**Table S1:**
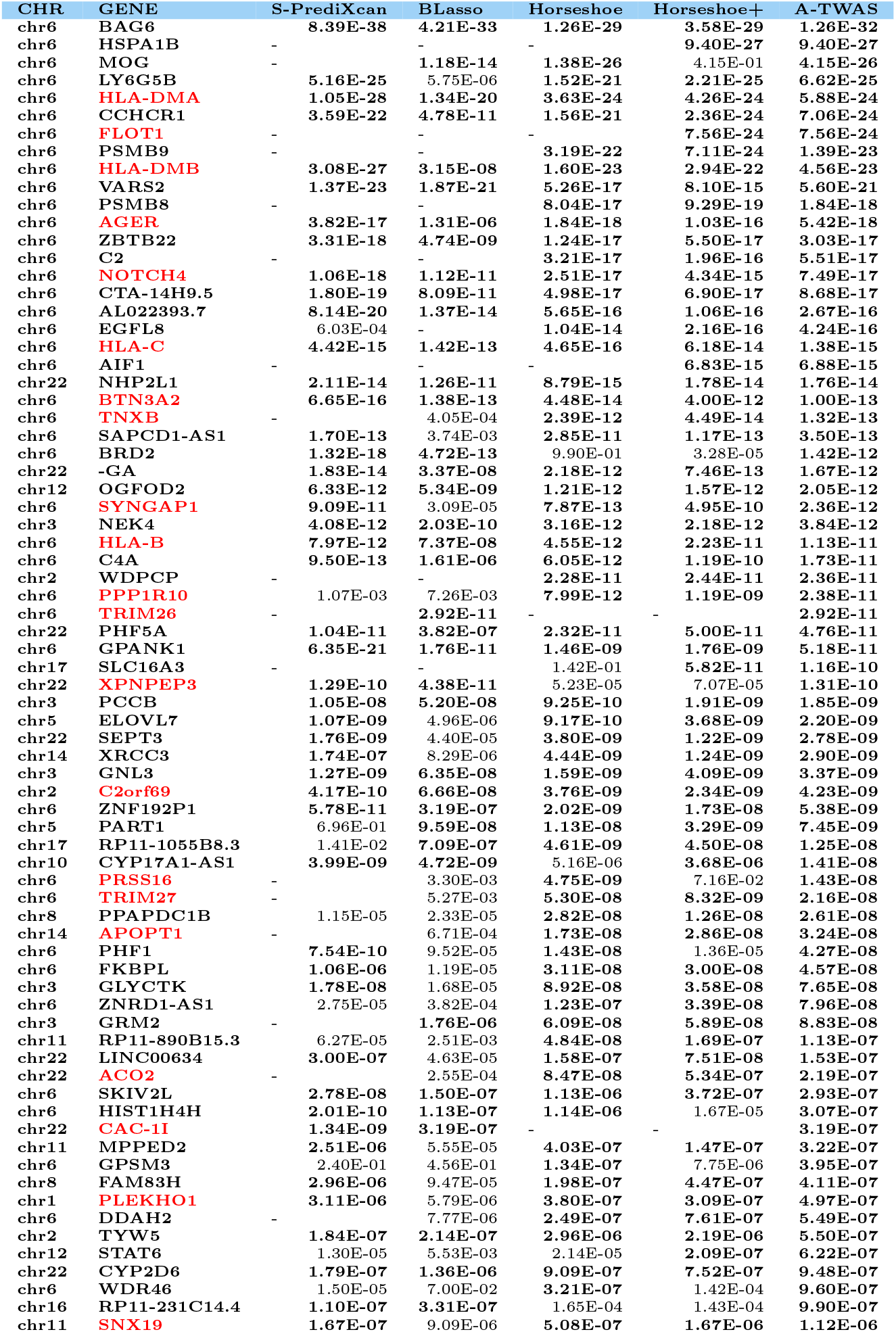
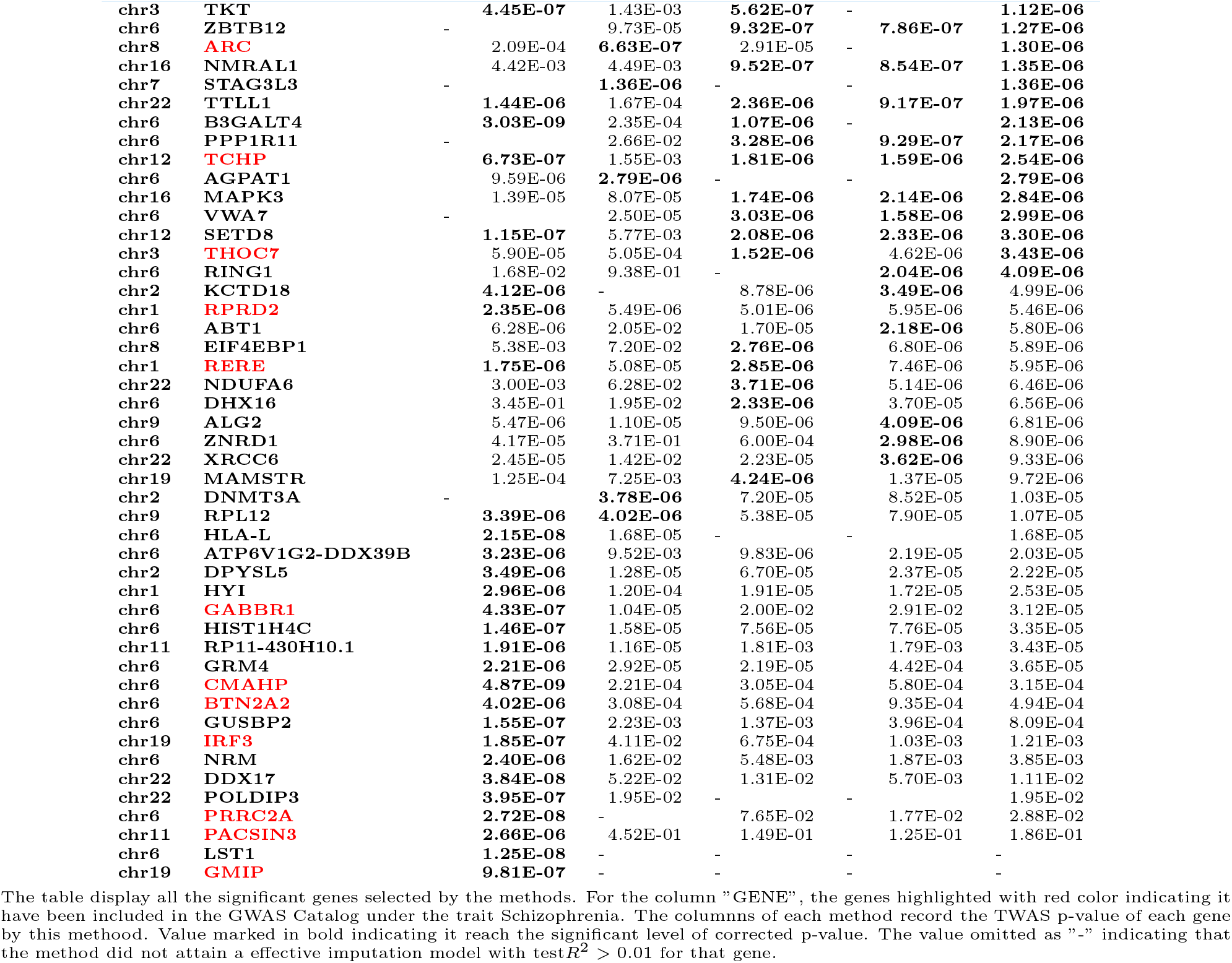
Significant Genes with TWAS p-value less than Bonferroni corrected significance level for Schizophrenia.

**Table S2:** Significant Genes with TWAS p-value less than Bonferroni corrected significance level for Obsessive-Compulsive Symptoms.

| CHR | GENE | S-PrediXcan | BLasso | Horseshoe | Horseshoe+ | A-TWAS |
| --- | --- | --- | --- | --- | --- | --- |
| chr6 | TPMT | - | - | **4.20E-06** | **4.24E-06** | **4.22E-06** |
| chr10 | ACTA2 | 7.84E-01 | 7.18E-01 | **1.02E-30** | 1.69E-01 | **3.06E-30** |
| chr3 | PARP15 | - | 3.17E-01 | **6.49E-09** | **6.16E-09** | **9.48E-09** |
| chr5 | MXD3 | 1.74E-01 | 1.93E-01 | **2.10E-13** | 2.65E-01 | **6.31E-13** |
| chr17 | SCO1 | 9.67E-01 | 9.00E-01 | **1.50E-06** | 7.54E-01 | **4.50E-06** |

The table display all the significant genes selected by the methods. The columns of each method record the TWAS p-value of each gene by this method. Value marked in bold indicating it reach the significant level of corrected p-value. The value omitted as "-" indicating that the method did not attain a effective imputation model with test $R^2 > 0.01$ for that gene.

## Notes

### Competing Interest Statement

The authors have declared no competing interest.

### Summary of Updates

Modified the manuscript template (removed the specific journal words)

